# MutationExplorer- a webserver for mutation of proteins and 3D visualization of energetic impacts

**DOI:** 10.1101/2023.03.23.533926

**Authors:** Michelle Philipp, Christopher W. Moth, Nikola Ristic, Johanna K.S. Tiemann, Florian Seufert, Aleksandra Panfilova, Jens Meiler, Peter W. Hildebrand, Amelie Stein, Daniel Wiegreffe, René Staritzbichler

**Author notes:** Authors contributed equally (Michelle Philipp, Christopher W. Moth, Nikola Ristic).

## Abstract

The possible effects of mutations on stability and function of a protein can only be understood in the context of protein 3D structure. The MutationExplorer webserver maps sequence changes onto protein structures and allows users to study variation by inputting sequence changes. As the user enters variants, the 3D model evolves, and estimated changes in energy are highlighted. In addition to a basic per-residue input format, MutationExplorer can also upload an entire replacement sequence. Previously the purview of desktop applications, such an upload can back-mutate PDB structures to wildtype sequence in a single step. Another supported variation source is human single nucelotide polymorphisms (SNPs), genomic coordinates input in VCF format.

Structures are flexibly colorable, not only by energetic differences, but also by hydrophobicity, sequence conservation, or other biochemical profiling. Coloring by interface score reveals mutation impacts on binding surfaces.

MutationExplorer strives for efficiency in user experience. For example, we have prepared 45,000 PDB depositions for instant retrieval and initial display. All modeling steps are performed by Rosetta. Visualizations leverage MDsrv/Mol*. MutationExplorer is available at: http://proteinformatics.org/mutation_explorer/

---

Mutations are essential to the persistence of life itself, enabling evolution of species in response to environmental pressures, exploited in protein engineering. Mutations can also cause challenging diseases. Throughout the biological sciences, we encounter perhaps *the* fundamental biophysical question: *How* does a mutation (or set of mutations) contribute to an observed or desired phenotype?

Specific flavors of this question include the search for explanations of phenotype variations, bacterial drug resistance, or human genetic disorders - to name only a few [1]. Many questions remain unanswered around cancer tissue, where evolution towards cell proliferation is so accelerated. In cancer, the molecular impact of mutation demands particular attention to understand disease etiology and progression.

Sequencing technologies have advanced rapidly in recent decades. As a result, over 50,000 organisms have been sequenced since 2018. Next generation sequencing (NGS) has reduced the cost of a complete human genome to less than US$1,000 [2] and today the Genome Aggregation Database (gnomAD 3.1.2) includes over 76,000 complete human genomes and over 125,000 exomes (sequenced protein-coding regions) [3]. The most frequently observed changes in proteins are missense mutations, where one amino acid is substituted by another [4]. Except for proline, all amino acids in a peptide chain share the same repeating backbone bonds atoms: -N-C*α*-C-. Thus, downstream effects of missense mutations arise from biochemical changes introduced by new side chain(s). In cases of side chains which bring great energetic strain, it can be hypothesized that a protein is destabilized, perhaps to the point that it can no longer fold correctly.

In cancer, NGS increasingly informs risk, diagnosis, prognosis, and therapeutic strategies [5]. Even though NGS is not frequently employed for routine laboratory and clinical diagnostics (as only special indications are directly diagnosed by NGS), the trend towards increased patient sequencing will continue, given cost reductions in sequencing and advances in personalized medicine.

While sequencing can reveal, even predict, risks of phenotype effects or disease, as of today, sequencing alone cannot explain the molecular mechanisms by which missense mutations drive changes in protein function. Analyzing these mechanisms requires attention to protein 3D structural context at atomic resolution. Indeed, amino acids quite separated in sequence can interact closely in a folded protein.

Here we present MutationExplorer webserver, which unites the advances in gene sequencing with advances in protein structure determination, and cements them with state-of-the-art computational methods and 3D graphics visualization. For any 3D protein model, MutationExplorer allows both the fluid exploration of mutation sets and a more systematic treatment of mutations through sequence uploads. Supported uploads include data from sequencing experiments, multiple sequence alignments, or reference sequences. On the server’s back end, Rosetta software [6] performs the mutations and calculates per-residue energies. On the web front end, mutated PDBs are displayed with the Mol* viewer [7]. Coloring scales reflect the differences in energy or hydrophobicity of the original compared to mutated proteins.

Through this flexible architecture, MutationExplorer integrates sequence and sequence variation with the exploding availability of 3D protein structures (both experimentally determined and modelled).

The Protein Data Bank (PDB)^3^ stores coordinates and related information on experimentally determined structures. However, most protein sequences are not directly represented in the PDB. Template-based modelling has long been available, and DeepMind’s AlphaFold has now leveraged machine learning to model the full length of most transcripts [8]. This opens the door to new modeling approaches which take protein structure as an input for further analysis. In particular, MutationExplorer benefits greatly from AlphaFold, as our webserver can now reveal predicted mutation energetics for a far wider array of sequence inputs.

## Methods and description

### Rosetta

The Rosetta software suite performs all computational modelling steps in MUTATIONEXPLORER. Rosetta debuted from David Baker’s group as a tool for structure prediction. Over the years, Rosetta has been extensively reworked and extended to encompass a variety of tools for protein design, docking, and other applications [9, 6]. Rosetta incorporates both physics and knowledge-based energy potentials, the latter potentials derived from principles of statistical physics and gleaned from analysis of high resolution structures in the PDB.

When predicting structures from sequence, Rosetta executes a full Monte Carlo conformational search for the given sequence. Many protein design protocols alternate between the design stage where the backbone conformation is held fixed, while a Monte Carlo algorithm focuses on replacement of the varying side chains, and a stage where the conformation of the newly designed sequence is explored. The design step of exchanging amino acids is accompanied by an overall side chain energy optimization to allow the local environment to adjust to the new arrival. Thus, prior to the design step, a pre-minimization is required. Without the pre-minimization mathematical artifacts are more likely to arise simply because the substitution and optimization could otherwise be driven by differences between the starting structure and the minimum determined by the Rosetta energy function. The pre-minimization typically moves the coordinates by only fractions of an Ångström, well within the error of experimental detection.

For the side chain optimization Rosetta employs a rotamer library gleaned from prior exhaustive conformational analysis of the PDB. Compared to gradient-based optimization on the rugged landscape of a vast conformational space, this rotameric approach drastically reduces the search space and automatically filters out many unfavorable side chain conformations from consideration.

Rosetta writes per-residue energies into each PDB output file. MutationExplorer writes these values into the B-factor column of the output file, allowing coloring of the entire protein molecule by its per residue energy or hydrophobicity - either in absolute values or as differences. That means, that for each coloring scheme an individual PDB file is created.

### Interface score

One of the visualization options shows the estimated effect of the mutation on binding energies in protein-protein complexes. To obtain these estimates, the InterfaceAnalyzer Rosetta mover [10] is called through PyRosetta [11] to calculate per residue binding energies of mutated and wild type structures. The difference (MUT - WT) is calculated for every residue. InterfaceAnalyzer calculates the binding energy by scoring the input structure twice: first, as a complex, and second, after moving the two sides of the interface away from each other, exposing the interface. The Rosetta energy scores of the unbound state are then subtracted from the scores of the bound one. Both the bound (input) and unbound structures have their side chains optimized prior to scoring. Since side chain optimization is stochastic, the mover is run 3 times, and the median score is taken for every residue.

### RaSP

RaSP is a new deep-learning-based tool that rapidly estimates protein stability changes. RaSP predictions strongly correlate to scores from Rosetta calculations which demand longer compute times [12]. With RaSP, MutationExplorer presents the user with a quick initial estimation of a mutation’s (de)stabilizing effect, without having to wait for the longer full minimization process. The RaSP tool consists of two linearly linked networks. A self-supervised 3D convolutional neural network that has learned representations of protein structures is followed by a fully connected neural network that maps these internal representations to Rosetta protein stability changes. RaSP is optimized for accuracy in the range [-1,7] kcal/mol, which is most relevant for revealing the loss-of-stability mutations underlying many diseases [13].

### Mol* viewer

Structure predictions are visualized in MutationExplorer with an adapted and extended version of Mol* [7], the successor of the NGL Viewer [14]. The extensions of Mol* from MDsrv [15, 16] have been incorporated into MutationExplorer so that sequence alignments are integratively displayed near structure visualizations. During development we were in close contact with the maintainers of Mol*, and we are incorporating our extensions into the Mol* main code branch. MutationExplorer structure visualization supports analysis in several complementary ways. After modeling the mutations, MutationExplorer highlights structural changes with a color gradient. Alternately, through the 1D sequence alignment control, the user can easily highlight and analyze the modified regions of the structure. In order to make introduced mutations easier to examine, the alignment view from MDsrv was integrated and its capabilities were extended. In MutationExplorer, this view now shows all the sequences of mutated versions of a protein and highlights the mutated loci in each sequence. Additionally, we developed a mutation highlighting approach which is active in the 3D structure visualization. Each mutated locus can be highlighted in a semi-transparent sphere, reducing search time and helping users to orient themselves as they rotate and translate the structure. The MutationExplorer back end conveys not only per-residue energetic changes to the user interface, but also hydrophobicity and sequence conservation metrics. The expert can select any of these values for coloring the mutated protein. MutationExplorer uses an adaptable color scale to visualize these differences so that the expert can tailor interpretation of impact to the various values in context of the protein of interest. When the mouse cursor is hovered over any amino acid, a panel appears which details the various computed values for the underlying residue. Clicking on a single amino acid selects it, and zooms in to the selected region. The side chain is displayed as well, to support regional visual inspection. From RaSP, every amino acid position is supported by an estimate of the global structure impact, should that position be mutated. The predicted values are shown by pressing the Ctrl key and left-clicking on the amino acid.

### Sequencing data pipeline (VCF input)

For human variants provided in GRCh38 genomic coordinates [17], chromosome, position, and alt alleles may be input in Variant Call Format (VCF) [18].

We employ the Ensembl Variant Effect Predictor (VEP) [19] to analyze the VCF file and return impacted Ensembl transcript IDs and amino acid variants. The VEP often returns IDs for computationally predicted splice variants which have not been experimentally verified. We filter to retain only Ensembl transcripts which cross-reference to Swiss-Prot curated, canonical UniProt transcript IDs [20]. Finally, MutationExplorer loads AlphaFold models [8] by UniProt ID and launches the analysis tools, as if mutations were input at protein level.

### Webserver

The MutationExplorer webserver is written in Python using the Flask framework with additional Javascript and is freely available for all users.

### AlignMe

alignme is a software package and webserver for detecting similarities between proteins too low to be detected on the sequence level using standard methods [21]. alignme is optimized for membrane proteins, but not limited to them. alignme offers a link on its result page that allows users to send their results to MutationExplorer for 3D visualization, providing the deepest insight into an alignment.

We have made available this same Javascript interface for other sequence alignment servers to forward their results for 3D visualisation to MutationExplorer. A link to MutationExplorer can be easily integrated with a few lines of code. Further details are in the FAQs of MutationExplorer.

### Usage

There are two ways to define mutations for a structure when using the web page provided by the MutationExplorer.

### The Upload Structure and Mutations option

is the way to go if the user wants to upload their own file or use a structure from a database such as the PDB or AlphaFold.

First, a 3D protein structure is selected in three different ways: by providing the file itself, by the definition of a PDB ID, or by the definition of an AlphaFold ID. Chains can be removed by clicking the Filter Structure button to clean up the structure. Additionally, the user can choose to minimize structures, calculate RASP predictions, and provide an email address to be notified when calculations are complete.

The mutations are defined in three different ways in the second step: by manual definition, by sequence alignment, and by definition of a target sequence. Manual definition is done by specifying which amino acid to switch at which position with a syntax like: ([chain]:[residue number][target mutation], example: “A:12G,B:134T”) If a sequence alignment is provided, the user simply selects the first sequence in the alignment to match to which chain in the structure. The definition of a target sequence is done either by uploading a fasta file, by writing out the target sequence, or by providing a UniProt ID for the chain that is to be modified. When this is done, the structure is ready to be mutated and the results will be calculated.

### The VCF upload

Alternately, a user can upload human SNPs, sequencing data in VCF format. For this second option, missense variants are obtained from Variant Effect Predictor output, and an AlphaFold model is selected automatically. In addition to the file upload, the user only has to decide on the minimization, the rasp calculation and whether or not to provide an e-mail address.

### Via the alignme server

It is also possible to import data from an alignme session. After the calculation of the results on the alignme web server, it is possible to export the result to the MutationExplorer. Selecting the export button on the server imports the clustal file from the calculation into MutationExplorer. The user just needs to select which sequence from the clustal file should be the base and target sequence, and provide the actual structural file for the sequences. This can be done by providing the ID of a PDB or AlphaFold structure. Alternatively, the file can be uploaded. Additionally, the user can choose whether the structures should be minimized and whether RASP predictions should be calculated.

### Navigating variants

MutationExplorer maintains a tree-structured list of previously explored variants. A user is free to return to any model in this tree, and define further mutations (which are added to the growing tree). The workflow is sketched in Figure 2. Newly created mutations are highlighted by spheres in the 3D view, and in bright-red in the sequence view.

**Figure 1.**
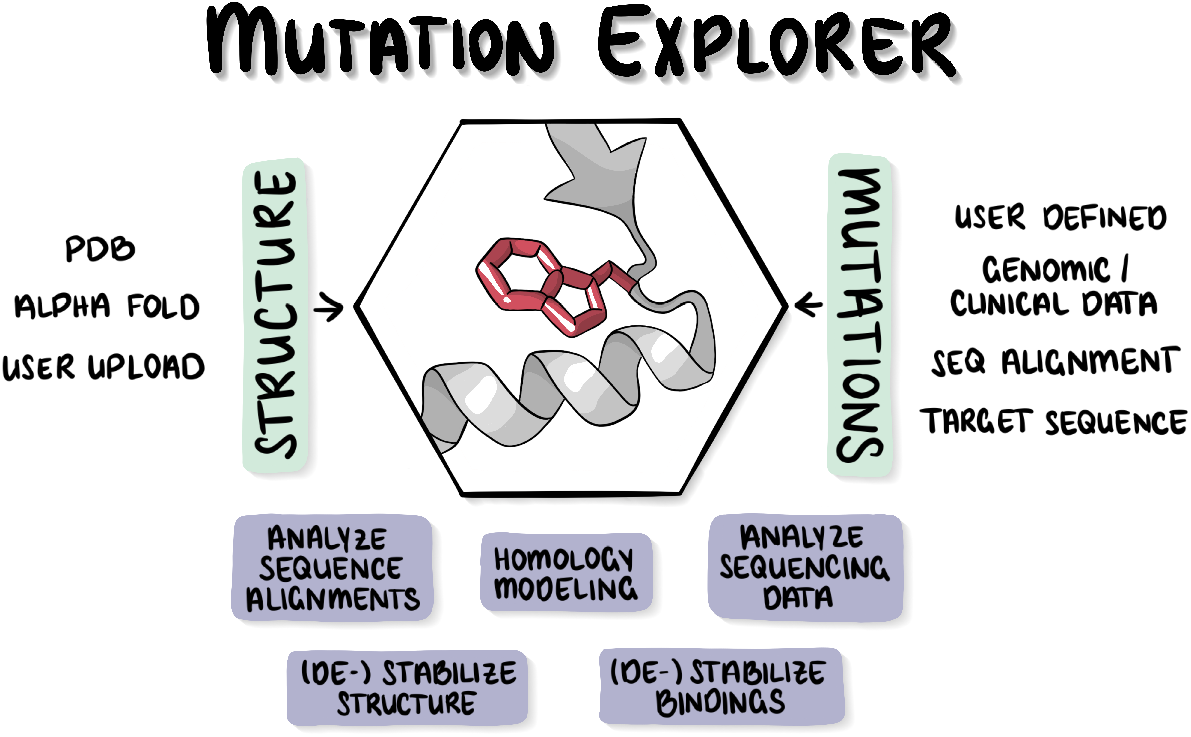
Graphical abstract. MutationExplorer integrates protein 3D structure and sequence variation with Rosetta energetics and presents results as a flexible and robust website.

**Figure 2.**
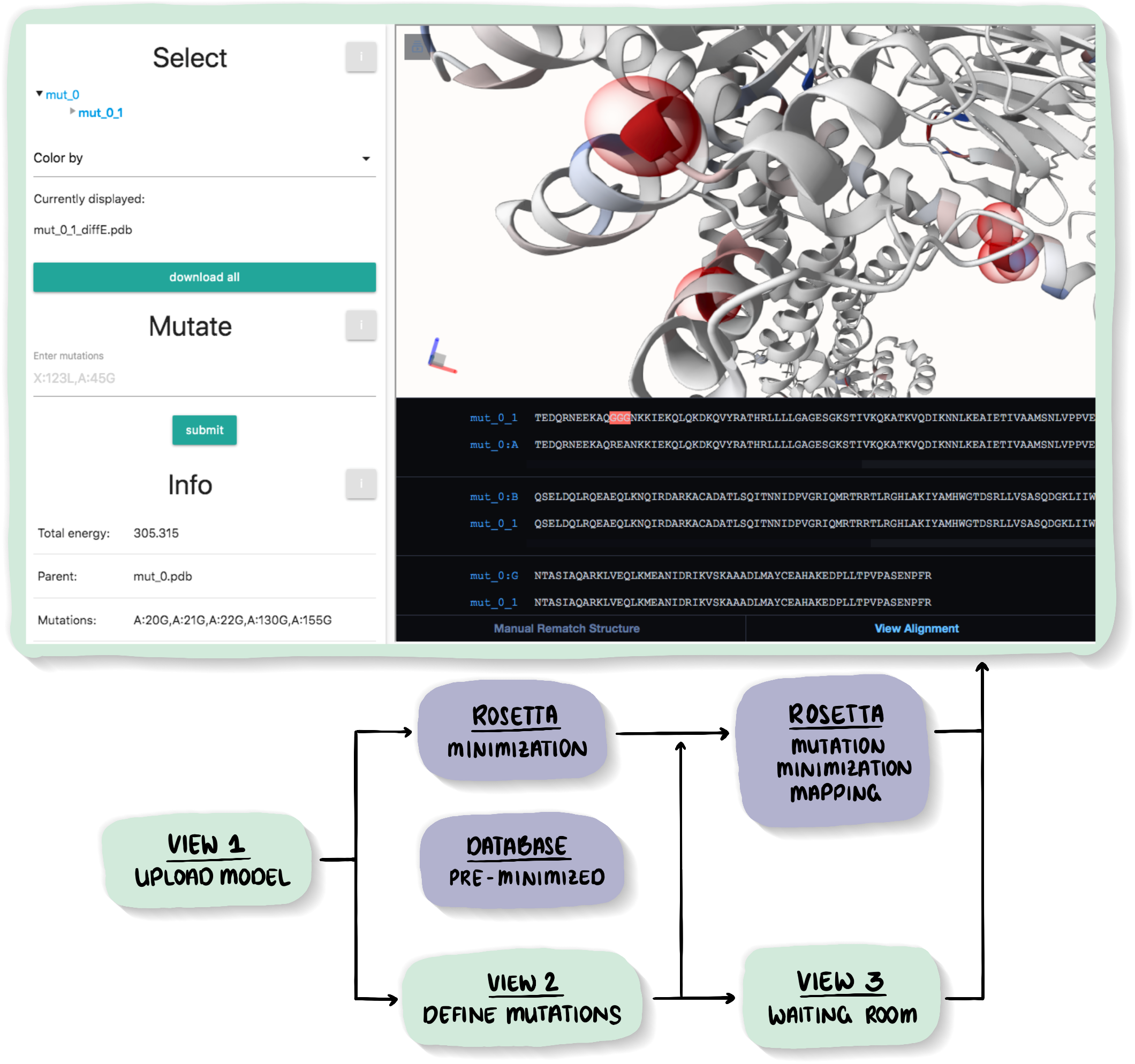
At the top, the result page is shown above a bubbled workflow schematic for the “upload structure and mutations” mode of the MutationExplorer server is shown. Below is a schematic of the (internal) workflow for structure and mutation preparation. There, the schematic *user steps* (the light green VIEW bubbles) are HTML pages displayed in the browser. Shown in slate purple bubbles are the *Rosetta modeling steps* that are executed in the background on the server. The user first uploads (or selects) a protein structure or model. Second, mutations are selected. Then a wäiting room” page is displayed while the mutations are performed. The result screen (top large rectangle, light green border) contains both the 3D viewer with the model of the mutated protein and below, linked to the 3D viewer, the sequence viewer shows the alignment. The same mutations are highlighted in red in both viewers. Hovering/clicking on a residue in either viewer highlights the residue in both. Additionally, a panel with details about the selected residue opens, including the energy/hydrophobicity value of the residue. Left mouse click adds an all atom licorice representation in the vicinity of the selected residue. A ‘Ctrl - left mouse button’ click opens a window with a fast estimate of the energy mutations at the given position will cause, calculated using RaSP [12]. The left section of the window has a selection tree of all created models, rooted at the original protein (‘mut_0’) and branching to all subsequent mutations. “Color by” choices include absolute or relative energies, as well as hydrophobicity. The user can continue exploring mutations by selecting a model and defining new mutations. Additionally, there is an information section, displaying the total energy of the cur8rently displayed variant, the parent from which the current model was derived together with the mutations that were performed on the parent. Finally, the user can adjust the color scale and toggle on/off the spheres highlighting the mutated positions.

### Coloring

All scores (energy, hydrophobicity, sequence conservation and interface binding) are available for coloration. For each score type, the user can switch colors between wildtype scores, mutation scores, and their differences.

Informed by this residue-wise coloring flexibility, users can quickly hypothesize mechanistic impacts of mutation for entire molecular systems or intermolecular binding interfaces.

### Navigation via sequence or 3D structure

A sequence alignment window is displayed beneath the structure window. Clicking on a residue in one window highlights it in the other, and vice versa.

### Visualization

Hovering reveals information about each residue. Left clicking displays side chain atoms. The “Change Visualization” button at the very bottom replaces the default sequence window with an interactive fine-tuner for color scales and overlaid spheres. The “View Alignment” button at bottom left re-displays the sequence window.

Additionally, explanations of the current color scheme are shown in an information panel. For example, sequence conservation colorings indicate the severity of an amino acid substitution using a range from white (no mutation) to light blue for mutations within amino acid groups to dark blue for mutations across groups. Sequence gaps are shown in red.

For any color scheme, a color gradient bar provides a key to the score range. Changing the bar’s flanking numeric values adjusts the displayed hue range.

### Mutation energy preview

Whenever RaSP calculations are included at launch (the default), the RaSP estimates of energetic effect are available via a ctrl-left click on any amino acid residue.

### Continuous mutation

From the results page you are free to add new variants to the current model. Previously entered variants automatically advance to subsequent model generations.

### Energy optimization

Global and thorough energy optimization before protein mutation is essential for high-quality results. Otherwise, our experience is that Rosetta will find mathematically “better” side chain conformations which are only distracting noise arising from the many small discontinuities between deposited atom coordinates and the Rosetta computational framework. Proper pre-optimization helps ensure that reported energetic differences are more directly related to the mutation itself. However, minimizing proteins, especially larger ones, is a time consuming step. MutationExplorer offers three kind of minimizations. First, we provide a database of pre-minimized proteins from the PDB. Both backbone and side chain minimizations were performed for these. For proteins not contained in the database, the server offers either a short or a long side chain optimization, using the *fixbb* application. Alternatively, the user can upload their own minimized models. Command lines are available in the Tutorial section on the website.

### A precalculated database

To build the pre-minimized database, we used a list of culled, non-redundant PDB identifiers from the PISCES server [22]. For the proteins contained in the list, we performed backbone minimization using the *relax* application of Rosetta, followed by a side chain optimization using the *fixbb* application. This yielded a database of pre-minimized structures for speeding users towards mutant exploration, currently containing 45,000 models.

Since the calculations for the minimization can be very time-consuming, the results for PDB and AlphaFold structures, which have not yet been optimized in advance, are added to the database so that the optimization does not have to be performed repeatedly. This is also done for the RaSP tool models.

### Limitations

Currently, MutationExplorer has no potentials targeted for membrane proteins available. Especially for residues facing the membrane, a special score-function and preparation is desired. Other residues, in particular those outside of the membrane can be investigated with MutationExplorer without limitations. Moreover, some mutations are generally challenging for Rosetta, foremost those from or to proline. Mutations from glycine may require conformational adaptation beyond the protocols used here. Mutations to glycine may cause proteins to be more flexible due to its larger torsion space. Ligands present in the PDB will be kept in rigid conformation, custom ligands will be removed. For advanced ligand handling, see [23]. MutationExplorer cannot handle very large structures, such as CryoEM structures available only in CIF format. Partial AlphaFold models for transcripts with more than 2,700 amino acids are not supported. We also caution that for large structures our default protocols for minimizations might be insufficient. Users should definitely minimize these on their local computer before upload. For this purpose, we recommend using the corresponding tutorial on Rosetta energy minimization via the command line, which can be found on our website. The fundamental challenge is the limited sampling of individual Monte-Carlo runs. A more reliable strategy for minimization is to create an ensemble of minimized models.

## Examples

The examples described below can be found on the server in the corresponding section in the main menu. In addition, there are more examples on the website under the Applications section of the tutorial, such as modeling multiple conformations of a protein and back mutation of PDB structures to wild type.

### RTEL1 VCF input example

As of this writing, 248 mutations in Telomere elongation helicase 1 (RTEL1) are classified as either “Pathogenic” or “Likely pathogenic” in the Clinvar database [24]. Of these, 13 are missense mutations. No experimental structure for RTEL1 is presently deposited in the PDB. To find structural trends in pathogenicity, we uploaded the 13 Clinvar SNPs to MutationExplorer in VCF format. Following our Rosetta based long-minimization and mutation protocols, MutationExplorer displays the AlphaFold model for canonical transcript Q9NZ71-1. 11 residues are highlighted in the red 3D structure and sequence viewers. Two of these Clinvar variants (Chr20:63662544 A*>*G and Chr20:63695619 G*>*A) do not impact the canonical uniprot transcript, and thus are not depicted. MUTATIONEXPLORER’s 11 red spheres depict all variants in obviously well-structured (high confidence) stretches of the AlphaFold model. Many of these variants involve mutations to or from proline, and MutationExplorer generally reports large energetic differences for these, consistent with general intuition around such mutations. Indeed, several fall at secondary structure transitional points near the ends of alpha helices. Perhaps more surprisingly, our Rosetta protocol calculated quite large energetic impacts (*>* 25 R.E.U.s) for the M492I [25] and I699M [26], both implicated in severe dyskeratosis congenita. More subtle energetic changes are also reported, reminding us that energetic analysis by itself, while necessary, is often insufficient. As example, the impact of V516L was computationally analyzed with an energetics-free model which based instead on 3D clustering patterns of pathogenic compared to benign variants [27]. In that study V516L clustered strongly with pathogenic variants, and was hypothesized to subtly disrupt DNA binding by adjacent surface residues. Link to example: proteinformatics.org/mutation_explorer/examples/rtel1/

### The GPCR autoproteolysis inducing domain

The GPCR autoproteolysis inducing (GAIN) domain is an extracellular domain of adhesion G-Protein coupled receptors, mediating receptor activation by releasing a autoproteolytically cleaved peptide from within itself [28, 29]. The GAIN domain therefore has two functions to serve - autoproteolysis and peptide release - which can be modulated by mutating the residues surrounding its cleavage site, the GPCR proteolytic site (GPS). In the rat ADGRL1 GAIN domain (PDB ID: 4DLQ), a conserved Trp residue, W804, has been demonstrated to abolish cleavage upon mutation [28]. This is reflected in substantial increases in energy for the W804S mutation in MutationExplorer - with a similar effect when mutating another conserved Trp, and W815S - by removing the bulky side chain. In the GAIN domain, two flexible regions termed “flaps” have been shown to mediate solvent-exposure of the GPS [29]. Point-mutations disrupting interaction of i.e. Flap 2 within residues 805-810, e.g. by Y806A, also result in energy increases not limited to the flap, but also in the interacting helix, hinting at a targeted disruption of the regions interactions, likely affecting overall GAIN domain dynamics. This example demonstrates the ability of MutationExplorer to predict the impact of point mutations on protein regions. Link to example: proteinformatics.org/mutation_explorer/examples/agpcr_demo/

## Availability

MutationExplorer joins our series of web services available at our platform http://proteinformatics.org which we aspire to support and maintain long-term: ProteinPrompt [30], MDCiao [31], Voronoia [32], Sierra Platinum Service[33], alignme [21]. MutationExplorer is freely available to all users online without any login requirement at: http://proteinformatics.org/mutation_explorer The MDSRV implementation on which MutationExplorer is build is available at: http://proteinformatics.org/mdsrv-web The interface for alignment servers is available at: https://github.com/starbeachlab/explorer_interface Issues with the website can be reported at: https://github.com/starbeachlab/MutationExplorerServer/issues

## Acknowledgements

We would like to express our gratitude to the team of developers and maintainers of Mol*, especially Alexander Rose, for many valuable remarks. The authors gratefully acknowledge the GWK support for funding this project by providing computing time through the Center for Information Services and HPC (ZIH) at TU Dresden.

## Declarations

The authors declare no conflict of interest.

## Footnotes

3 https://www.wwpdb.org

